# Dynamic plant height QTL revealed in maize through remote sensing phenotyping using a high-throughput unmanned aerial vehicle (UAV)

**DOI:** 10.1101/369884

**Authors:** Xiaqing Wang, Ruyang Zhang, Liang Han, Hao Yang, Wei Song, Xiaolei Liu, Xuan Sun, Meijie Luo, Kuan Chen, Yunxia Zhang, Guijun Yang, Yanxin Zhao, Jiuran Zhao

## Abstract

Plant height is the key factor for plant architecture, biomass and yield in maize (*Zea mays*). In this study, plant height was investigated using unmanned aerial vehicle high-throughput phenotypic platforms (UAV-HTPPs) for maize diversity inbred lines at four important growth stages. Using an automated pipeline, we extracted accurate plant heights. We found that in temperate regions, from sowing to the jointing period, the growth rate for temperate maize was faster than tropical maize. However, from jointing to flowering stage, tropical maize maintained a vigorous growth state, and finally resulted in a taller plant than temperate lines. Genome-wide association study for temperate, tropical and both groups identified a total of 238 quantitative trait locus (QTLs) for the 16 plant height related traits over four growth periods. And, we found that plant height at different stages were controlled by different genes, for example, *PIN1* controlled plant height at the early stage and *PIN11* at the flowering stages. In this study, the plant height data collected by the UAV-HTTPs were credible and the genetic mapping power is high, indicating that the application of this UAV-HTTPs into the study of plant height will have great prospects.

**Highlight:** We used UAV-based sensing platform to investigate plant height over 4 growth stages for different maize populations, and detected numbers of reliable QTLs using GWAS.

## Introduction

Maize (*Zea mays*) was domesticated from Balsas teosinte (*Zea mays* subspecies *parviglumis*) in southwestern Mexico around 9,000 y BP (van Heerwaarden et al., 2011). Subsequently, maize has been continuously improved by humans, and the most important improvements were spread from the tropical region to the temperate region, which can be called adaptation (Liu et al., 2015). The adaptation process allowed maize to be widely cultivated worldwide and become the largest production food crop in the world (http://faostat3.fao.org/compare/E). However, the world population is soaring and the demand for food is also increasing. It has been reported that the world’s grain demand must meet a target of 70% increase by 2050 (Tester and Langridge, 2010). Therefore, corn, the largest grain, has become particularly important in safeguarding world food security.

Maize yield is highly complex and is affected by many factors, among which plant height is a particularly important factor because it not only affects the lodging resistance, but also biomass and yield (Salas Fernandez 2009). In the first Green Revolution, with the successful application of the semi-dwarf genes (*rht1*; *sd1*) in wheat and rice, the crop yields increased dramatically (Peng et al., 1999; Khush et al., 2001; Sasaki et al., 2002). Plant height was so important that people have made unremitting efforts to exploring its genetic mechanism. So far, there were plenty of quantitative trait loci (QTLs) identified for maize plant height using a diversity of genetic populations (Peiffer et al., 2014; Yang et al., 2014; Dell’Acqua et al., 2015; Zhou et al., 2016; Pan et al., 2017). Some of these genes were cloned, such as *an1*, *dwarf3*, *dwarf8*, *dwarf9* and *br2*, which were mainly involved in the synthesis and transportation of gibberellin and auxin (Winkler and Freeling, 1994; Bensen et al., 1995; Winkler et al., 1995; Fujioka et al., 1988; Xing et al., 2012).

Maize plant height showed different characteristics during the whole growth period, especially in the important growth stages, such as the seeding, jointing, flowering and mature stages (Abendroth et al., 2011). Usually, maize grows slowly in the seedling stage, fast in the jointing stage, then gradually slower in the grouting stage, and stops in the milky stage (Zhang et al., 2012). However, for a long time, researchers have often investigated the plant height at the mature stage to obtain the final height, leading to a lack of systemic understanding of the entire plant height development process and the genetic factors of its genetic development mechanism. Furthermore, the workload of manual measurement also contributed to plant height typically only being investigated at one growth stage.

Manually investigating plant height is a laborious and time-consuming task. Since plants are tall at maturity, errors are unavoidable in the measurement process and the accuracy of the data will be affected. In recent years, with the development of artificial intelligence, a series of high-throughput automated phenotypic systems have been developed. At present, indoor platform systems are widely used for dissecting phenotypic traits in which environmental effects are minimized (Yang et al., 2014; Zhang et al., 2016; Al-Tamimi et al., 2017); however, field high-throughput platforms have much fewer applications within the complex environment that farmers routinely experience (Crain et al., 2016; Liang et al., 2018). Compared with indoor platforms, the development of field high-throughput platforms requires high flexibility and a large payload (Araus and Cairns, 2014). Thanks to the advance in remote sensing, aeronautics and high-performance computing development, some field-based high-throughput phenotypic platforms (HTPPs) have been developed (Araus and Cairns, 2014). For example, the Australian Plant Phenomics Facility (http://www.plantphenomics.org/hrppc/capabilities/technology), and ground-based HTPPs used for wheat, cotton, sorghum and maize, which can determine the canopy height, reflectance, temperature, plant height, biomass and so on (Andrade-Sanchez et al., 2013; Holman 2016; Duan et al., 2017; Liang et al., 2018). However, these field-based HTPPs have very few applications in genetic improvement, especially for genetic mapping.

To better understand the dynamic plant height mechanism, we investigated the plant height through four growth periods with an unmanned aerial vehicle (UAV) system for maize diversity inbred lines, which covers wide genetic diversity and is widely used in maize genetic research (Yang et al., 2011; Yang et al., 2014; Liu et al., 2017). Through this design, we hope to explore more plant height characteristics with the aid of the high-throughput UAV and data processing procedures, and then dissect the genetic basis for plant height for different maize groups at different stages.

## Materials and methods

### Plant materials and experiment design

The maize natural population used in this study was a subset of Yang (Yang et al., 2010), consisting of 117 temperate lines and 135 tropical lines, which had a high-density genotype of 1.25 million single nucleotide polymorphism (SNPs) with minor allele frequency (MAF) more than 0.05 (Liu et al., 2017). The population was sown on 15 May 2017 at Xiao Tang Shan, Changping, Beijing National Precision Agriculture Research Center of China (115°E, 40°N). The land plots were flat, with uniform soil fertility. There was a row length of 2 m, including eight plants, and each line included three rows. Row-to-row distance was set as 65 cm. Phenotypic data collection with UAV was carried out on 8 June, 29 June, 11 July and 3 August 2017, days with clear sky and no wind (Table 1; Fig. 1A). On the same days, the height of 44 randomly selected plants was manually measured with a ruler.

**Fig. 1.**
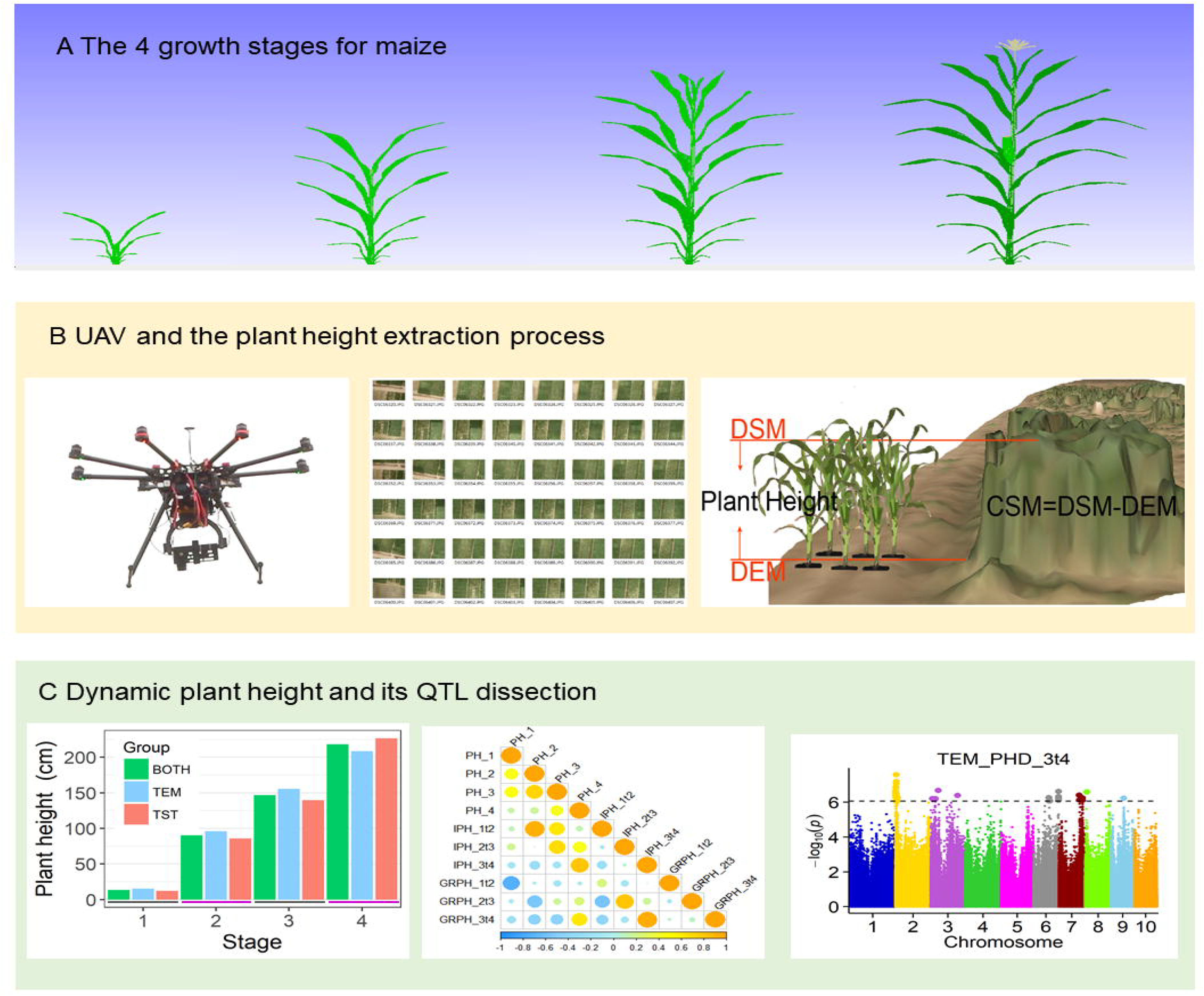
Field high-throughput phenotyping for plant height. A, digital designed graphs for the maize plants during the four stages. B, the UAV equipment and the plant height extraction process. The main process contained the image collection by the UAV, then divided the pictures into the mosaic plots, and extracted plant height based on the formula (CSM=DSM-DEM). C, Dynamic plant height and QTL dissection. The whole procedure included trait variation and correlation analysis, as well as GWAS.

**Table 1.** The investigation date for plant height

| **Flight** | **Date** | **DAS<sup>a</sup>** | **DAFD<sup>b</sup>** | **Development stage** | **Description** |
| --- | --- | --- | --- | --- | --- |
| 1 | 8 June 2017 | 24 | — | V5 | seeding stage |
| 2 | 29 June 2017 | 45 | 21 | V10 | jointing stage |
| 3 | 11 July 2017 | 57 | 12 | V12 | trumpet |
| 4 | 3 August 2017 | 80 | 23 | R | flowering stage |

Note:
a: DAS means days after sowing
b: DAFD means days after the closest former date

### Platform and image acquisition

An Octocopter UAV (DJI Spreading Wings S1000) platform was used to collect a set of aerial images across four flights (Fig. 1B). A 20.2-megapixel digital camera (Cyber-shot DSC-QX100) was mounted on the UAV to acquire the images by means of a global positioning system (GPS) and inertial navigation unit system. In each flight, the same flight plan was followed with 80% forward overlap and 75% side overlap at an altitude of approximately 40–60 m, depending on the sun situation. The flight routes were programmed into the UAV software to automatically generate efficient flight paths for UAV. Each flight speed was set to 6 m/s. International Standards Organization sensitivity and shutter speed were set to automatic, and the focal length was fixed at 10.4 mm. The flight time was within 15 min. Sixteen ground control points (GCPs), measured using millimeter-accuracy differential GPS (South Surveying & Mapping Instrument Co., Ltd., China), were evenly distributed in the field to obtain an accurate geographical reference from multiple dates.

### Plant height image data extraction and verification

Digital surface models (DSMs) and orthomosaics were produced from images shot by UAV with GCPs using the structure-from-motion software Agisoft PhotoScan 1.3 (Agisoft LLC, St. Petersburg, Russia) (Fig. 1B). This process included feature point matching, dense point cloud generation, product output, etc. A digital elevation model (DEM) (i.e., a non-vegetation ground model) was constructed from the first set of aerial images collected 24 days after sowing by the local polynomial interpolation method. Crop surface models (CSMs) were calculated by subtracting the DSM at different plant growth stages from the DEM (Fig. 1B, Hoffmeister et al., 2010; Bendig et al., 2013; Hoffmeister et al., 2013). The CSM includes a raster dataset that mixes the soil and plant pixels. Many studies have shown that extracting plant height directly from CSM results in underestimation (Bendig et al., 2015; Holman et al., 2016; Watanabe et al., 2017). Segmenting plants from soils using the excess green index proposed by Woebbecke et al. (1995) was a necessary measure for the above extraction. Kriging spatial interpolation and maximum adjacent pixel methods were performed on CSMs to remove the soil background, and the maximum of interpolation was taken as the representative value of plant height at the plot scale. To assess the accuracy of plant height extraction from UAV, 44 maize plants were randomly selected to manually measure plant height at the second, third and fourth timepoints of plant growth. A linear regression model was applied with multiple dates using R v. 3.2.4 statistical software.

### Plant height variation between temperate and tropical maize

A total of 252 maize inbred lines, consisting of 117 temperate lines and 135 tropical lines, were used in this study. Plant heights were evaluated at four different growth stages, and a total of 16 plant height related traits were calculated, including 4 absolute plant height traits (PH), 3 incremental plant height difference (IPH), 3 growth rates of plant height (GRPH), 3 daily incremental plant height difference (DIPH) and 3 daily growth rates of plant height (DGRPH). The PH represents the absolute plant height at each timepoint. The IPH represents the difference between the adjacent timepoints, e.g. IPH_1t2 equals PH_2 minus PH_1. The GRPH was calculated as the ratio of IPH divided by the former plant height, e.g. GRPH_1t2 equals IPH_1t2 divided by PH_1. The DIPH was the IPH divided by the total days between the adjacent timepoints. Finally, the DGRPH was calculated as the GRPH divided by the total days between the adjacent timepoints. The phenotypic distribution and graphs were implemented in the R v. 3.2.4 statistical software.

### Association analysis for plant height

Genome-wide association study (GWAS) was carried out in temperate (TEM), tropical (TST) maize and both of the two e population (BOTH). Genotype data quality control was performed separately, with 1,141,328, 1,110,483 and 1,227,441 SNPs remaining for TEM, TST and BOTH groups, respectively. We used 16 plant-height-related traits in the GWAS program, including PH, IPH, GRPH, DIPH and DGRPH traits for the three groups. Combined with phenotypes and genotypes, the FarmCPU model in the MVP software package, which iteratively uses fixed and random effect model, was used for association tests in TEM and TST groups with only kinship considered (Al-Tamimi et al., 2016; Liu et al., 2016). For the BOTH population, the top five principal components were added in FarmCPU model to control false positives, which may be caused by population stratification and the non-genetic effect (Al-Tamimi et al., 2016; Liu et al., 2016). The adjusted Bonferroni method (i.e., P ≤ 1/N, where N is the total number of genome-wide SNPs) was used as the global P value cutoff to declare significance of SNPs associated with a given trait. The P values were 8.76e−7, 9.0e−7 and 8.14e−7 for the TEM, TST and BOTH populations, respectively. QTL intervals were calculated as the upstream and downstream 100kb for each significant SNP (Deng et al., 2017). Any SNP in the QTL interval with the lowest *P* value was considered as the peak SNP.

We searched the genes in each QTL according to the physical position of each gene in maizeGDB (https://www.maizegdb.org/). Gene annotations were based on both maizeGDB and InterProScan database (http://www.ebi.ac.uk/interpro/scan.html). The gene expression profiles were also from maizeGDB.

## Results

### High-throughput digital plant height extraction and validation

To investigate the plant height of 117 temperate and 135 tropical maize inbred lines for the four stages, we used the unmanned aerial vehicle high-throughput phenotypic platform (UAV-HTTP) system to collect the image data. We carried out four flights during the whole development stage of maize during the seedling, jointing, trumpet and flowering periods (the V5, V10, V12 and R stages at 24, 45, 57 and 80 days after seeding, respectively). On each flight, the average flight altitude was 52.5m. A total of 559 original images were taken on four flights (Table 2). Using the self-developed automated data extraction process, we first filtered the original images, and retained 460 high-quality images, with an average of 115 images per flight. After the reconstruction of the orthomosaic model, the obtained image ground resolution was 1.15 cm/pixel. The DSM was constructed using the orthomosaic model output point cloud data. The average image accuracy of the DSM was 2.31 cm/pixel. The DEM was generated by interpolation of the DSM points located on the surface of the bare land. Finally, we obtained the CSM containing bare soil (DSM − DEM, Fig. 1B). Here, the tiny terrain at the bottom of the crop can be ignored because whole plant area was flatted by a farmland leveling machine. Therefore, CSM is equivalent to crop height. The average coefficient of variation (CV) of plant height gradually decreased from 53% to 11.6% among first three growth stages caused by the increasing heterogeneity of plant height. The mean crop height ranged from 9.6 to 253.4 cm among the four periods with an average growth rate of 4.06 cm/d (Table 2).

**Table 2.** Features for the extraction for the plant height using UAV-HTTP

| **Flight** | **Flight**<br>**Altitude**<br>**(m)** | **Original**<br>**images**<br>**quantity** | **Checked**<br>**images**<br>**quantity** | **Orthomosaic**<br>**Resolution**<br>**(cm/pixel)** | **Point**<br>**Density**<br>**(points/cm-2)** | **DSM**<br>**Resolution**<br>**(cm/pixel)** | **Min of**<br>**CSM**<br>**(cm)** | **Max of**<br>**CSM**<br>**(cm)** | **CV of**<br>**CSM**<br>**(%)** | **Mean of**<br>**CSM(cm)** |
| --- | --- | --- | --- | --- | --- | --- | --- | --- | --- | --- |
| 1 | 40 | 166 | 120 | 0.72 | 47.9 | 1.44 | 0 | 26 | 53 | 9.6 |
| 2 | 60 | 113 | 98 | 1.33 | 14.2 | 2.65 | 69 | 184 | 13.4 | 124.9 |
| 3 | 60 | 121 | 95 | 1.35 | 13.7 | 2.71 | 117 | 251 | 11.6 | 185.9 |
| 4 | 50 | 159 | 147 | 1.23 | 16.4 | 2.47 | 148 | 365 | 14.5 | 253.4 |

To verify the accuracy of the plant height data extracted using UAV-HTTP, 44 lines were randomly selected for plant height measurement by ruler at the same time as the 2nd, 3rd and 4th flight. A linear regression model was established for the UAV-HTTP data and ruler measurement data and the model correlation coefficient was very high (r^2^= 0.91), indicating that the data obtained by the UAV platform had high accuracy (Fig. 2).

**Fig. 2.**
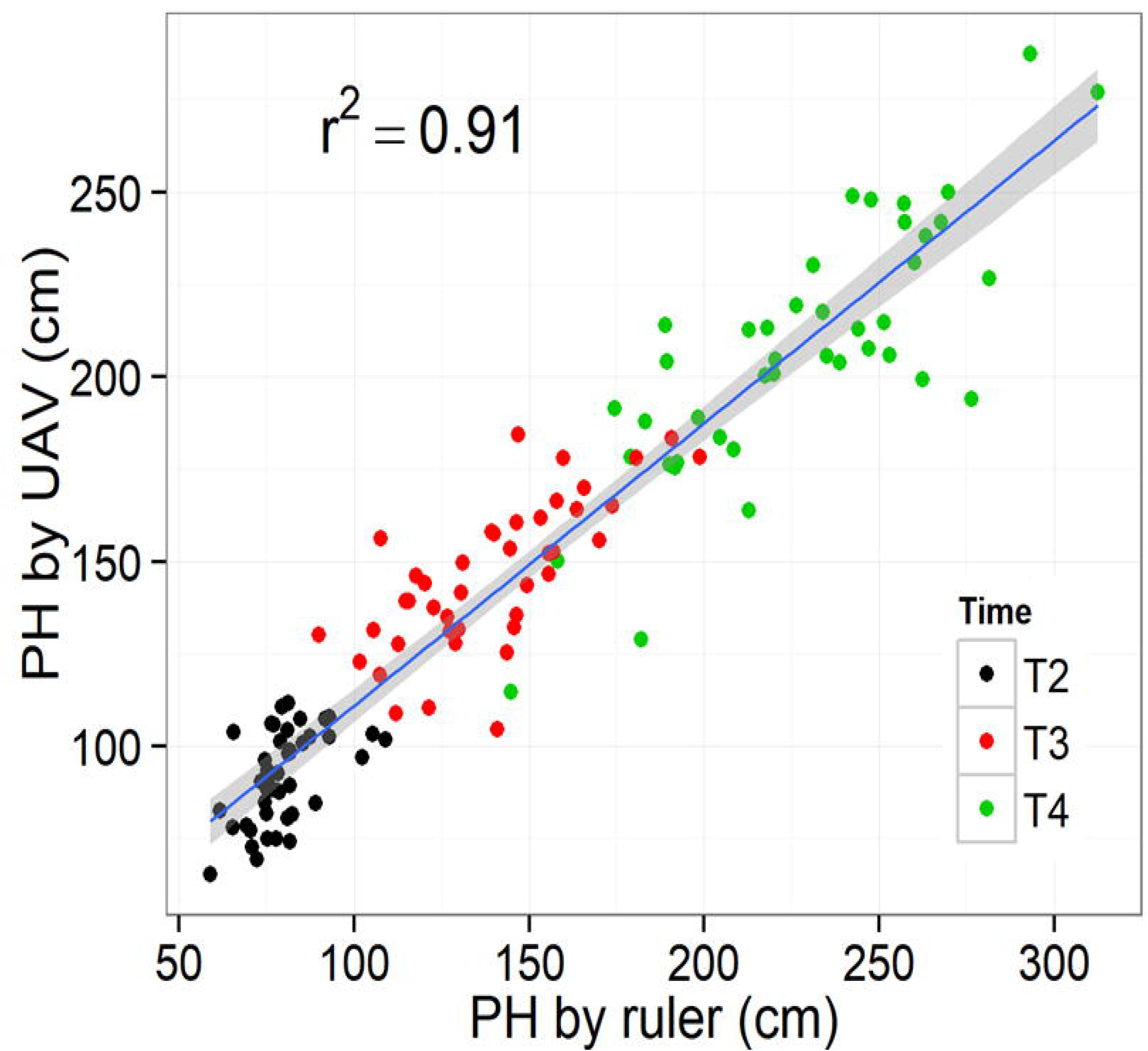
Linear relationship for plant height by UVA and manual measurement by ruler at three growth stages. The blue solid line represents the regression line, and the grey shadow represents the 99% confidence interval.

### Plant height varies greatly at different stages of development

Based on the accurate plant height data obtained by UAV-HTTP, we performed further analysis of the variation for maize plant height across the four growth stages among the three different groups (Table 3). For the BOTH group, the average PH were between 13.66 and 218.26 cm, from the first to the fourth flight (Fig. 1C). The DIPH values for the three adjacent periods were from 3.68 to 4.74 cm, with the maximum for 2t3 stages, and minimum for 3t4 stages. However, the DGRPH values varied from 0.02 to 0.31, with the maximum for 1t2 stages, and minimum for 3t4 stages. The inconsistence for DIPH and DGRPH indicate that growth rate was not positively correlate to incremental growth.

**Table 3.**
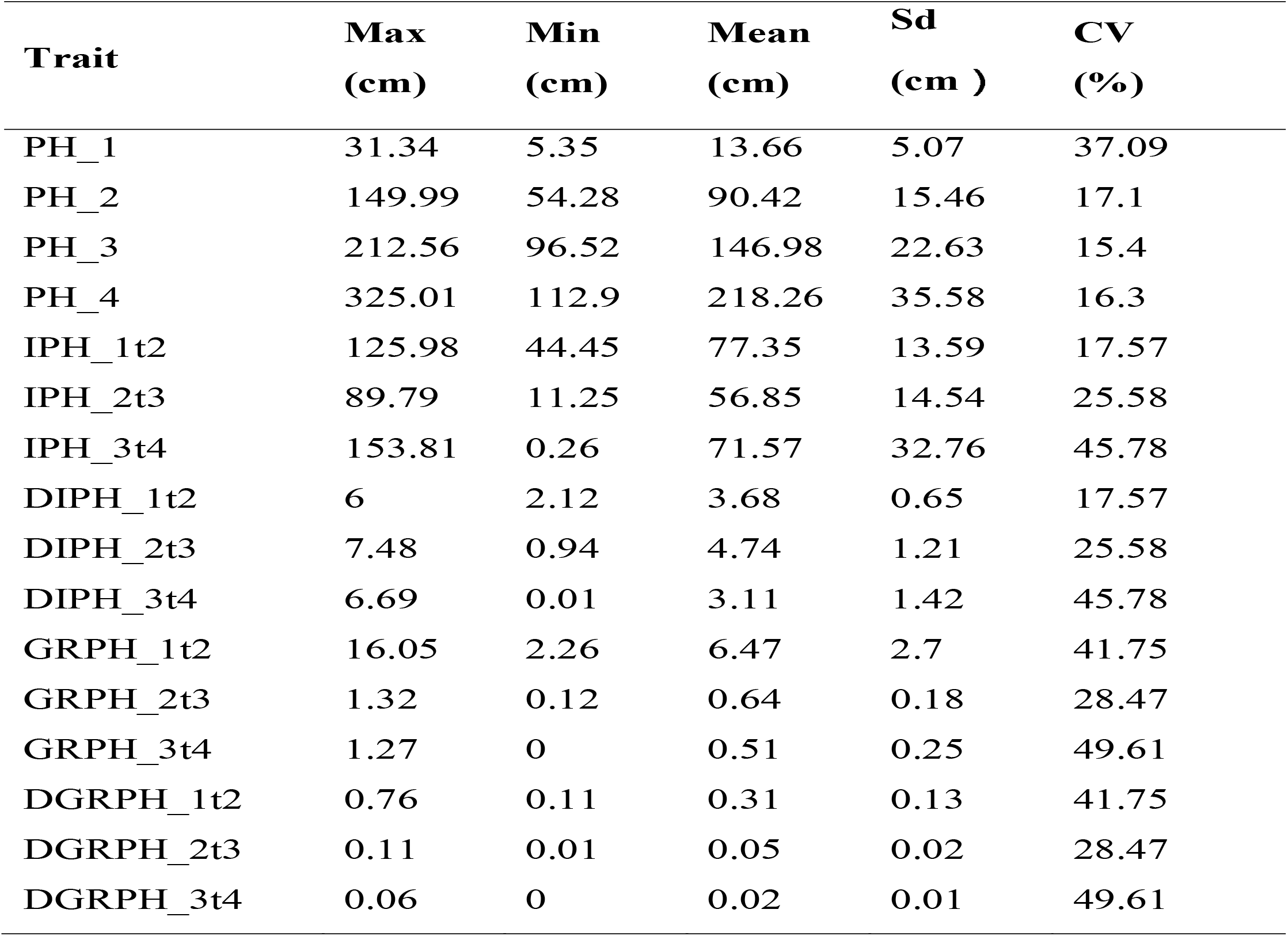
Statistic analysis for plant height variation for the whole population during four growth stages

| **Trait** | **Max**<br>**(cm)** | **Min**<br>**(cm)** | **Mean**<br>**(cm)** | **Sd**<br>**(cm )** | **CV**<br>**(%)** |
| --- | --- | --- | --- | --- | --- |
| PH_1 | 31.34 | 5.35 | 13.66 | 5.07 | 37.09 |
| PH_2 | 149.99 | 54.28 | 90.42 | 15.46 | 17.1 |
| PH_3 | 212.56 | 96.52 | 146.98 | 22.63 | 15.4 |
| PH_4 | 325.01 | 112.9 | 218.26 | 35.58 | 16.3 |
| IPH_1t2 | 125.98 | 44.45 | 77.35 | 13.59 | 17.57 |
| IPH_2t3 | 89.79 | 11.25 | 56.85 | 14.54 | 25.58 |
| IPH_3t4 | 153.81 | 0.26 | 71.57 | 32.76 | 45.78 |
| DIPH_1t2 | 6 | 2.12 | 3.68 | 0.65 | 17.57 |
| DIPH_2t3 | 7.48 | 0.94 | 4.74 | 1.21 | 25.58 |
| DIPH_3t4 | 6.69 | 0.01 | 3.11 | 1.42 | 45.78 |
| GRPH_1t2 | 16.05 | 2.26 | 6.47 | 2.7 | 41.75 |
| GRPH_2t3 | 1.32 | 0.12 | 0.64 | 0.18 | 28.47 |
| GRPH_3t4 | 1.27 | 0 | 0.51 | 0.25 | 49.61 |
| DGRPH_1t2 | 0.76 | 0.11 | 0.31 | 0.13 | 41.75 |
| DGRPH_2t3 | 0.11 | 0.01 | 0.05 | 0.02 | 28.47 |
| DGRPH_3t4 | 0.06 | 0 | 0.02 | 0.01 | 49.61 |

We conducted a correlation analysis for the BOTH group to reveal the relationship between the 16 traits at different plant stages (Fig. 3). A strong positive correlation was found between IPH and DIPH, GRPH and DGRPH. Second, correlations for PHs at different stages were also positively related, from 0.14 to 0.73. Third, the correlation between IPHs was weak ranging from −0.25 to 0.05. PHRs were also weakly related to each other, from −0.09 to 0.1. However, there was a positive correlation between IPH and PH, ranging from −0.28 to 0.94. The correlation between GRPH and PH was mainly negative, ranging from −0.73 to 0.62. In addition, the correlation between IPH and GR was relatively variable, ranging from −0.57 to 0.95.

**Fig. 3.**
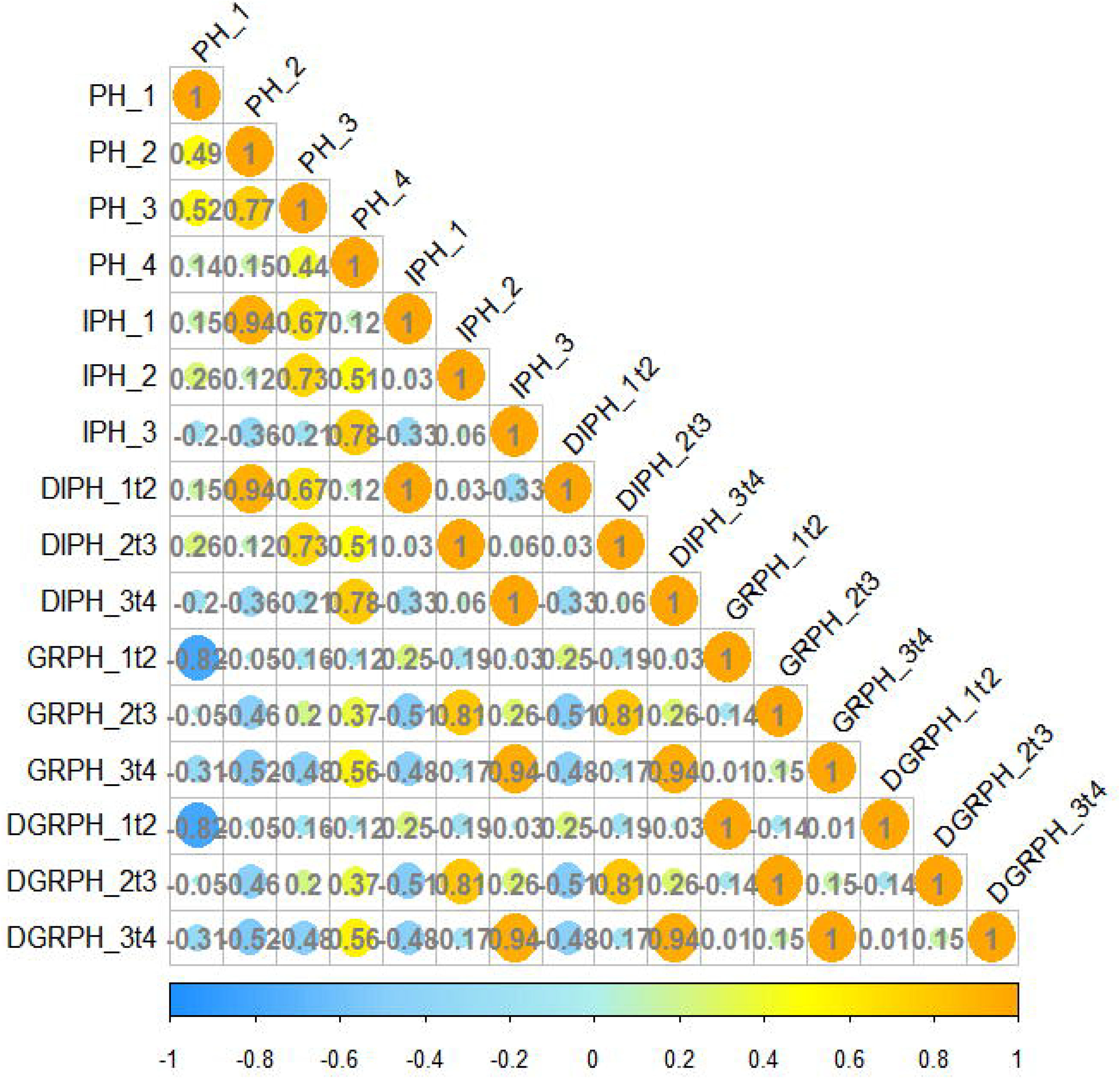
Correlation coefficient matrix among 16 plant-height-related traits. Yellow and blue indicate positive and negative correlations, respectively, and the size of the circle is proportional to the correlation coefficient. The number indicates the correlation coefficient.

As the wide diversity of the BOTH group, we divided the group into TEM and TST groups, and found the plant height between TEM and TST maize exhibited a significant difference at each growth stage (Fig. 4). From the first to the third flight (namely 1t2 and 2t3 stages) the DIPH values for TEM and TST were 3.87 vs. 3.52 cm and 4.97 vs. 4.53 cm, respectively, showing that TEM maize consistently grew faster than the TST maize. However, from the third to the fourth period, the TST grew faster than the TEM maize, and the DIPH for TEM and TST were 2.32 vs. 3.79 cm. More importantly, the most significant difference for the two groups were at 3t4 stage when most TEM lines were flowering, while most TST lines were still in vegetative growth.

**Fig. 4.**
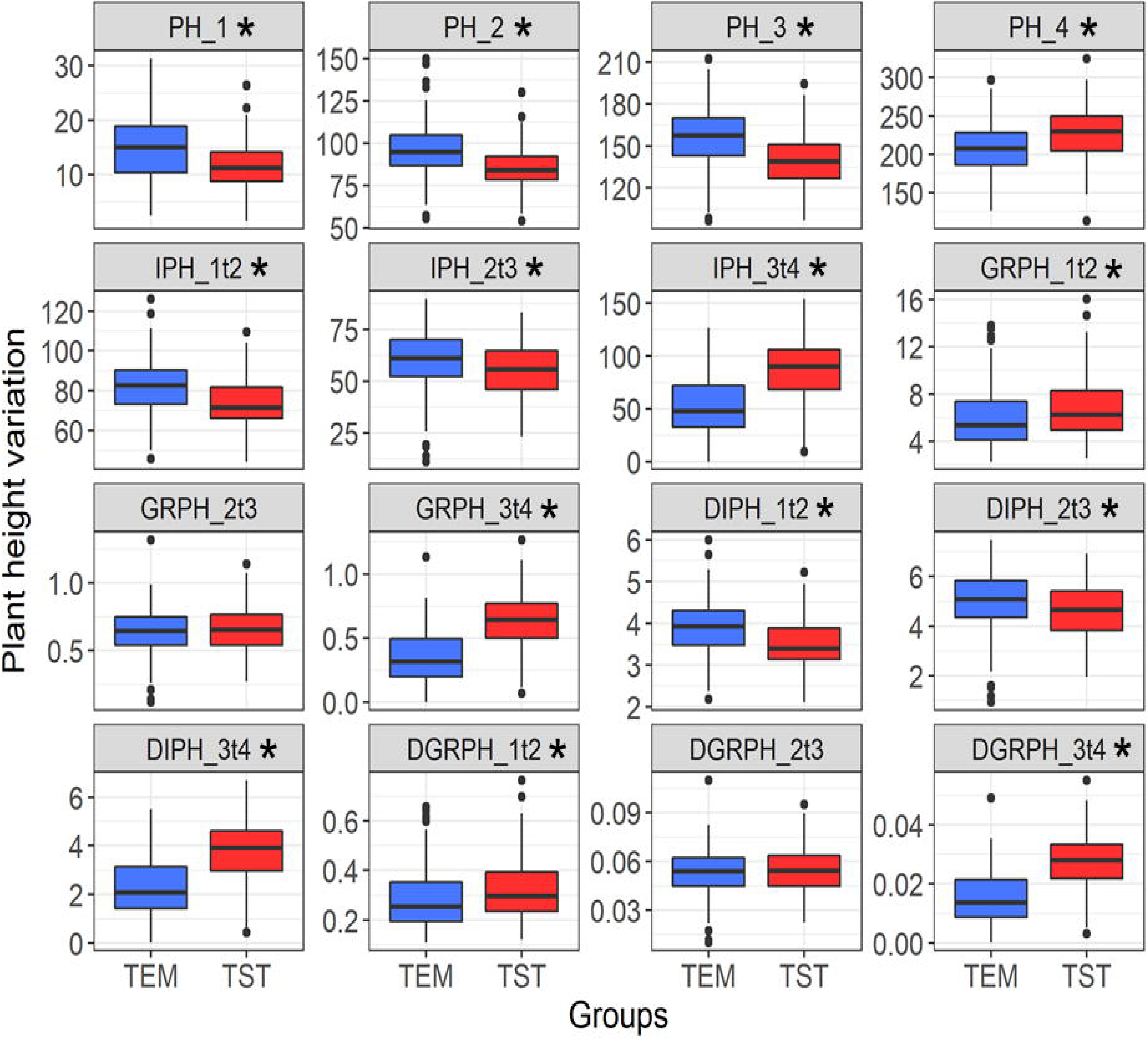
Plant height and its related trait variations between the TEM and TST populations at four growth stages. Blue and red represent the TEM and TST populations, respectively. The line in the box plots show the median value. Box edges represent the first and third quartiles, and the dots outside the whiskers represent the value over 1.5 × interquartile range. Stars means phenotypic distribution has significantly difference below 0.05.

### Genetic basis affecting the dynamic development of plant height

In view of the above-mentioned differences in plant height and related traits in different groups and at different stages of growth, we conducted GWAS for the 16 plant heigt related traits in the TEM, TST, and BOTH groups. A total of 238 QTLs were detected, covering 10 chromosomes of the maize genome (Data S1-S5; Fig. S1-S4). There were 38, 49, 50, 50 and 51 QTLs detected for PH, IPH, GRPH, DIPH and DGRPH traits, respectively.

To verify the accuracy of the QTL, we compared the previously reported QTLs and genes related to plant height and found that 45% of the QTLs overlapped with previous research (Peiffer et al., 2014; Yang et al., 2014; Dell’Acqua et al., 2015; Zhou et al., 2016; Pan et al., 2017). In addition, genes involving the GA and auxin pathway were also detected to be associated with plant height, such as *ARFTF4*, *D3*, *GA2OX8*, *KS3*, *PIN1* and *PIN11*, indicating that the QTL results of this study were highly reliable (Tudroszen et al., 1977; Winkler and Helentjaris,1995; Lo et al.,2008; Yamaguchi, 2008; Li et al., 2016; Weijers et al., 2018). Furthermore, 55% of QTLs were newly identified in the present study, including traits related to plant height and growth rate. Combining a large number of validated and new QTLs, we can discover the genetic basis affecting the dynamic development of plant height.

First, plant height at different stages was controlled by different QTLs. There were 6, 6, 2 and 24 loci detected for PH traits at the V5, V10, V12 and R stages, respectively (Fig.5; Data S1). More QTLs at the flowering stage were detected than at other stages. However, comparison of the QTLs for the four developmental stages did not show any overlapping regions, suggesting that plant height was controlled by different genes at different stages. For example, at the V5 stage, the gene *PIN1*, an auxin transport protein (Kumari et al., 2015), was detected near to the QTL of chr9: 3.23-3.43Mb for the TEM group. The expression profile of *PIN1* in B73 showed high expression in the early stem, indicating that the gene may involve in the early stages of development. At the R stage, the gene *PIN11* which is an auxin efflux carrier family protein (Tudroszen et al., 1977; Balzan et al., 2014), was detected in the QTL of chr2:192.33-192.53Mb for the TEM group. The expression profile of *PIN11* for B73 showed that the gene expressed highly in the SAM and internode, indicating that the gene was likely to have controlled plant height, especially in later development.

**Fig. 5.**
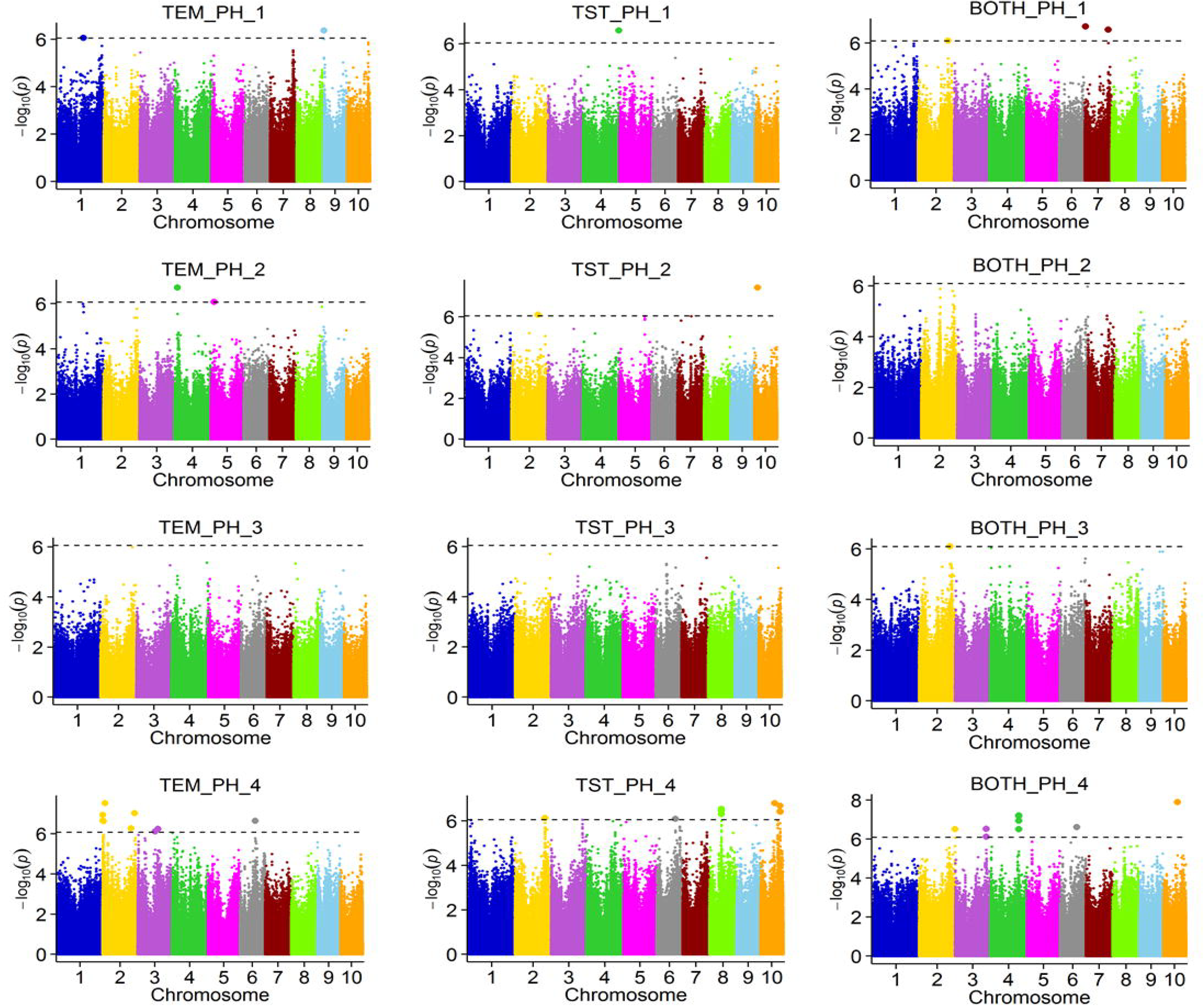
Genome-wide association study for the plant heights at the four stages among the TEM, TST and BOTH groups. Different colors represent different chromosomes. The dotted line is the threshold. SNPs above the threshold showed significant association ones.

Second, different QTLs controlled plant height in different groups. We found that few QTLs for the three populations overlapped in the same stage. For the PH trait across four stages, there were more QTLs detected in the other two groups than BOTH group except in the first stage (Fig. 6). The reason may be different GWAS model used for the three groups, and the model used for BOTH group was strict than the other two groups with population structure be considered. In addition, for the similar model of TEM and TST groups, the QTLs were still different, which may be caused by the allele frequencies. For example, the QTL chr2: 192.33-192.53 Mb (containing *PIN11*) was only detected in the TEM group at the R stage. The allele frequencies of the peak SNP (chr2.S_192432591, A/G) were 0.52/0.47 in the TEM group, 0.16/0.85 in the TST.

**Fig. 6.**
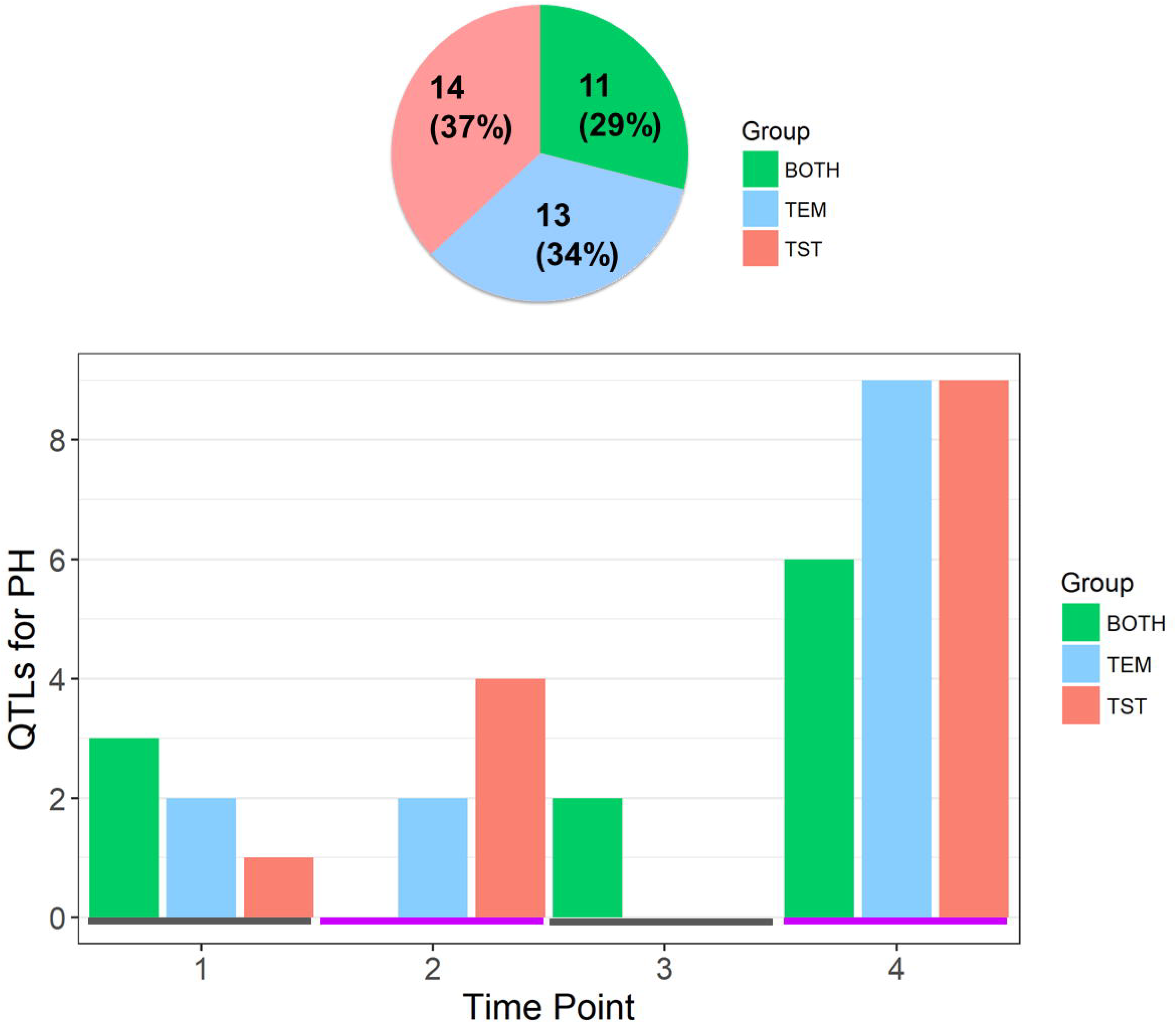
Number of QTL for plant height among the three groups (TEM, TST and BOTH) for four stages. A, the number and proportion of QTLs for the three groups. B, the QTLs for each group at each of the four stages.

Third, we have found considerable overlap between PH, IPH, GRPH, IDPH and DGRPH (Fig. 7; Fig. S5). Based on the correlations between the five class traits, and the results of co-localization of QTLs, we can obtain a systematic understanding of the genetic basis of traits. For example, the QTL region chr2:2.49-4.36 Mb was co-located by PH_4, IPH_3t4 and GRPH_3t4, which contained the auxin corresponding factor gene *ARFTF4* (Auxin response factor 4; Li et al., 2016), indicating that the plant height at the R stage was mainly contributed by the difference of the growth rate of V12 to R stages rather than other periods. These results indicate that the plant height surveyed at a specific stage was affected by many factors, such as IPH and the plant height at the former stage.

**Fig. 7.**
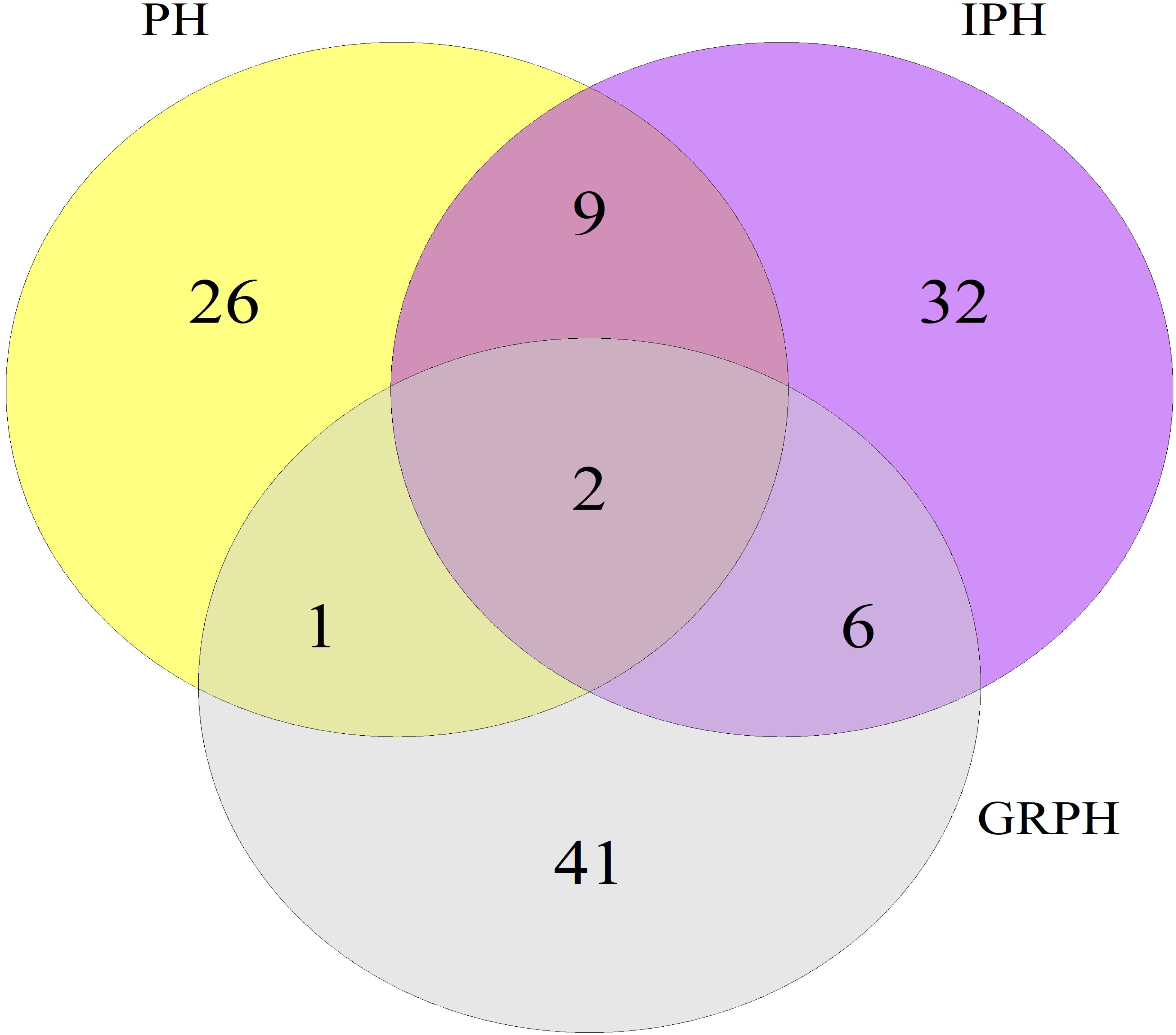
The venn graph of QTLs for PH, IPH and GRPH traits.

## Discussion

### High-throughput phenotyping platforms promote genetic research

Application of genetic improvement is the most effective way to increase crop yields. With the fast development of sequencing technology, genomic researches have recently been rapidly increasing; however, the phenotype has been facing bottleneck (Furbank and Tester, 2011). The development of HTPPs to obtain more phenotypes has been the focus of the fast development of genetics and breeding.

A series of indoor phenotypic platforms have recently been developed and applied into genetic researches (Chen et al., 2014; Yang et al., 2014; Zhang et al., 2016). The application of these high-throughput, automated phenotyping devices can greatly shorten the phenotypic investigation time, ensure the accuracy of the phenotype, and discover phenotypes that researchers cannot obtain by conventional techniques. More importantly, the traits discovered by the high throughput platform can identify some known genes as well as the novel loci, providing a valuable ability for gene identification.

Compared with indoor platforms, the development of field HTPPs will be much more complex because of the requirement for high flexibility and a large payload (Araus and Cairns, 2014). To date, UAV has been an excellent tool as field high-throughput techniques, and has achieved great success in the researches of wheat and cotton (Andrade-Sanchez et al., 2013; Holman et al., 2016). However, the applications for UAV in maize plant height research were very few. In this study, we applied the UAV platform to survey maize plant height in the fields and used the resultant accurate data for genetic mapping. A large number of reported and many novel QTLs were detected, showing the advantage of GWAS using the UAV-HTPPs in mining of plant height loci. The platform is likely to have a wide range of future applications and can be extended to more complex agronomic traits.

### Dynamic phenotype accelerates the dissection of the genetic basis of plant height

The determination of plant height variation depends on the in-depth investigation of phenotype. Currently, the survey of plant height typically takes place at the mature stage, which can obtain stable traits, but a lot of useful plant height information is likely to be missed. In this study, we monitored the plant height from the seeding stage to the flowering stage, through division into four periods. We found that GRPH of maize varies greatly at different stages of development, with the fastest in 1t2 stage, and slowest in 3t4 stage. Second, we found that TST maize grew slower and had a shorter plant height than the TEM maize from sowing to jointing stage. However, from the jointing to the flowering stages, TST maize had a faster growth rate, and finally resulted in a taller plant than TEM maize. Third, there were different genes regulating the plant height at different stages, some controlling early growth, some controlling mid-term and some controlling later stages. In this study we have detected 6, 6, 2 and 24 QTLs for the PH traits at V5, V10, V12 and R stages, but no common QTLs among the four stages. The results were consistent with Yan (Yan et al., 2003), who investigated plant heights in five periods and found that QTLs controlling plant heights were expressed differently in different periods. The above results indicate that if we assess the plant height over different growth stages, we will be able to identify more genes affecting plant height. Fourth, we found that a few regions can be co-localized by PH, IPH and GRPH. For example, we found that the co-localized QTLs controlling later IPH or GRPH were also detected in later PH traits and vice versa. This indicates that by dividing the plant height into several stages of growth, the key factors for the plant height can be better identified at specific stages. The dynamic phenotype enables us to have a clearer understanding of plant developmental processes. The usage of dynamic phenotypic data for mapping can identify more QTLs affecting the development of the trait, which is of great importance for the analysis of the genetic basis of traits and subsequent improvement of the trait.

## Acknowledgements

The authors are grateful to Dr. Jianbing Yan for providing the maize natural population. Thank Dr. Boxiang Xiao for the software of Maize Three-Dimensional Interactive Digital Design (PlantCAD-maize) which helps to design the digital maize plants.

## Author contributions

Y.X.Zhao, G.J.Y, and J.R.Z designed the experiment; R.Y.Z, X.S, K.C and Y.X.Zhang carried out the field experiment; G.J.Y and L.H conducted the UAV-HTTP. X.Q.W implement the statistical analysis and GWAS work. X.Q.W and Y.X.Zhao prepared the initial draft. X.L.L, M.J.L and W.S helped to modify the manuscript. J.R.Z and W.S provide the foundation support. All authors reviewed the manuscript.

## Abbreviations

BOTH: Both of temperate and tropical maize
CSMs: Crop surface models
CV: Coefficient of variation
DEM: Digital elevation model
DGRPH: Daily growth rate of plant height
DIPH: Daily incremental plant height
DSMs: Digital surface models
EGI: Excess Green Index
GCPs: Ground control points
GPS: Global positioning system
GRPH: Growth rate of plant height
IPH: Incremental plant height
MAF: Minor allele frequency
PH: Plant height
QTL: Quantitative trait locus
SNP: Single nucleotide polymorphism
TEM: Temperate maize
TST: Tropical maize
UVA: Unmanned aerial vehicle
UAV-HTPPs: Unmanned aerial vehicle high-throughput phenotypic platforms

